# Novel *Rickettsia* spp. in two common overwintering North American passerines

**DOI:** 10.1101/2022.06.18.496698

**Authors:** Daniel J. Becker, Allison Byrd, Tara M. Smiley, Mariana Fernandes Marques, Julissa Villegas Nunez, Katherine M. Talbott, Jonathan W. Atwell, Dmitriy V. Volokhov, Ellen D. Ketterson, Alex E. Jahn, Kerry L. Clark

## Abstract

American robins and dark-eyed juncos migrate across North America, but their contributions to arthropod-borne disease remain poorly characterized. We identified novel *Rickettsia* spp. in one wintering migrant per bird species related to bellii, transitional, and spotted fever group rickettsiae and suggest spring migration could disperse these pathogens hundreds-to-thousands of kilometers.

## Main text

Migratory birds play an important role in shaping risks of arthropod-borne diseases, given their ability to disperse pathogens and vectors to new wintering or breeding locations. For pathogens with high public health burdens, such as *Borrelia burgdorferi* and spotted fever group (SFG) rickettsiae, migratory birds can disperse millions of infected vectors annually (1). Many migratory birds are also competent for these pathogens and therefore can not only disperse pathogens and arthropods but also transmit infections to naïve vectors following migration (2).

American robins (*Turdus migratorius*) and dark-eyed juncos (*Junco hyemalis*) are important species to understand bird migration and arthropod-borne disease in North America. They are competent for some pathogens and can have high ectoparasite intensities (2,3). Both have diverse migratory behaviors, are widely distributed, and are common in suburban habitats (4,5), which could facilitate pathogen dispersal to areas of high human exposure. Yet the contribution of these common migratory bird species to arthropod-borne disease remains poorly understood.

Here, we sampled robins and juncos across North America for arthropod-borne pathogens of public health significance. We focused on bacterial pathogens for which these birds are known to be competent (*Borrelia* spp.) (2), for which detections have occurred in other passerines (*Rickettsia* spp.) (6), and that are rarely or never seen in birds (*Bartonella* spp., hemoplasmas).

Robins were sampled monthly in southern Indiana (2020–2021), while juncos were sampled in southern California (2006), Virginia’s Appalachian Mountains (2018, 2019), and northeastern Ohio (2019). California and Ohio were sampled in the breeding season, while Virginia was sampled in winter (7). We captured birds with mist nets, applied USGS bands, and collected blood stored in 96% ethanol or Longmire’s buffer at –20°C, on Whatman FTA cards, or frozen directly at –20°C. We also collected the first secondary feather for stable isotope analyses. Sampling was approved by the Indiana University IACUC (06-242, 18-028), Federal Bird Banding Permit 20261, and state permits. Table 1 shows sample sizes per site and month.

**Table 1.** Sample sizes for robins and juncos per month, year, and region. All samples were tested for *Borrelia* spp, *Rickettsia* spp., and *Bartonella* spp.; parentheticals indicate samples tested for hemoplasmas. Asterisks indicate the single *Rickettsia*-positive per bird species.

| Avian host | Date | Region | Number sampled |
| --- | --- | --- | --- |
| *Turdus migratorius* | 1/2020 | Southern Indiana | 21 (8) |
| *Turdus migratorius* | 2/2020 | Southern Indiana | 15 (8) |
| *Turdus migratorius* | 3/2020 | Southern Indiana | 77 (41) |
| *Turdus migratorius* | 4/2020 | Southern Indiana | 67 (32) |
| *Turdus migratorius* | 5/2020 | Southern Indiana | 8 (7) |
| *Turdus migratorius* | 9/2020 | Southern Indiana | 18 |
| *Turdus migratorius* | 10/2020 | Southern Indiana | 20 |
| *Turdus migratorius* | 11/2020 | Southern Indiana | 25 |
| *Turdus migratorius* | 12/2020 | Southern Indiana | 24* |
| *Turdus migratorius* | 1/2021 | Southern Indiana | 21 |
| *Turdus migratorius* | 2/2022 | Southern Indiana | 1 |
| *Turdus migratorius* | 3/2022 | Southern Indiana | 18 |
| *Turdus migratorius* | 4/2022 | Southern Indiana | 37 |
| *Turdus migratorius* | 7/2021 | Southern Indiana | 5 |
| *Turdus migratorius* | 9/2021 | Southern Indiana | 28 |
| *Turdus migratorius* | 10/2021 | Southern Indiana | 6 |
| *Junco hyemalis* | 2/2006 | Southern California | 25 |
| *Junco hyemalis* | 3/2006 | Southern California | 54 |
| *Junco hyemalis* | 4/2006 | Southern California | 32 |
| *Junco hyemalis* | 5/2006 | Southern California | 25 |
| *Junco hyemalis* | 6/2006 | Southern California | 24 |
| *Junco hyemalis* | 7/2006 | Southern California | 9 |
| *Junco hyemalis* | 7/2019 | Northeastern Ohio | 20 |
| *Junco hyemalis* | 11/2018 | Appalachian Mountains | 65* |
| *Junco hyemalis* | 11/2019 | Appalachian Mountains | 30 |

DNA was extracted from blood with Maxwell RSC Whole Blood DNA kits or Qiagen DNeasy 96 Blood and Tissue kits. We then used published PCR protocols to screen avian DNA for *Bartonella* spp. (partial *gltA* gene), *Borrelia* spp. (partial 16S rRNA gene), and *Rickettsia* spp. (23S-5S rRNA intergenic spacer [ITS]). A subset of robins was also tested for hemoplasmas (partial 16S rRNA gene). Table S1 provides PCR primers and thermocycler conditions.

Of 675 samples, we detected rickettsiae in one robin (0.26%, 1/391) and one junco (0.35%, 1/284), representing the first reports of rickettsiae in these bird species. No other target pathogens were detected (Table 1). Both sequences (GenBank accessions ON773823 and ON773824) shared only 82.61% partial identity of the 23S and 5S rRNA flanking sequences of their ITS to one another, indicating distinct rickettsiae. We used NCBI BLASTn to identify related rickettsiae 23S-5S rRNA ITS and type strain sequences, followed by MUSCLE for alignment and MrBayes for phylogenetics (10,000,000 generations, GTR+G+I) via NGPhylogeny.fr (8).

Both sequences had ≤91% identity to other *Rickettsia* spp. The robin sequence was related (83.85–91.01% identity with 91% sequence coverage) to uncultured rickettsiae from *Ixodes auritulus* in Argentina (MW824654), *Amblyomma americanum* and *Ixodes scapularis* in the USA (KJ796407 and KJ796403), humans in Ethiopia (MK693112) and India (OK077732–OK077742), and to *R. monacensis* (e.g., JQ796867), *R. tillamookensis* (CP060138), and *R. felis* (e.g., DQ139799). The junco sequence was only partially related (87.63–88.66% identity with 49% sequence coverage) to the above sequences, and the main internal part (146 bp) of this ITS sequence (without 23S and 5S rRNA flanking sequences) was unique with no identity to any rickettsiae ITS sequences in GenBank. These sequences thus belong to two novel but not yet cultured *Rickettsia* spp., and our 23S-5S ITS phylogeny suggested they are most similar to rickettsiae within the bellii group (BG), transitional group (TRG), and SFG (9) (Fig. 1A).

**Figure 1.**
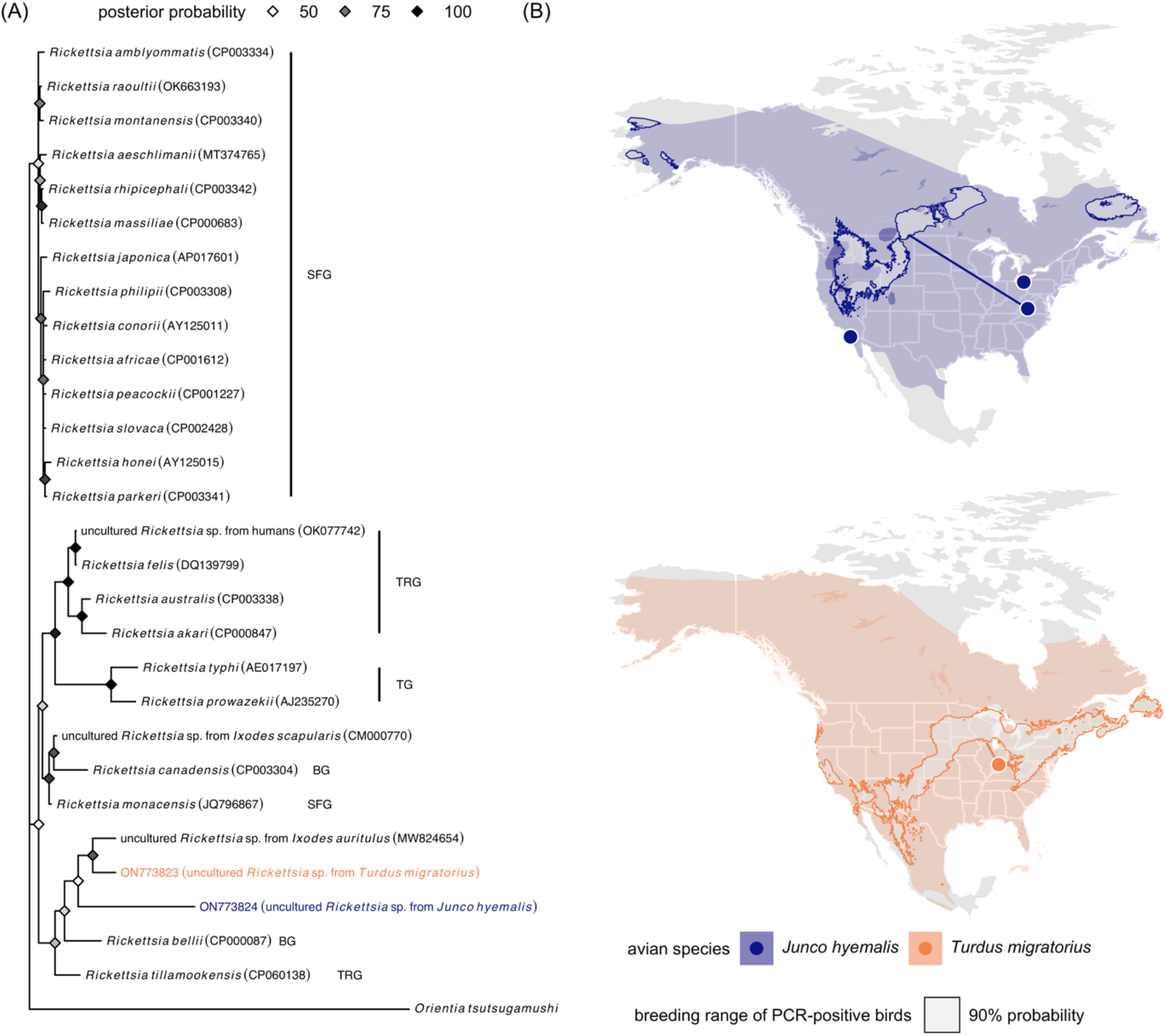
(A) Bayesian phylogeny of the novel *Rickettsia* spp. with closely related and reference rickettsiae sequences from GenBank, including those in the bellii group (BG), typhus group (TG), transitional group (TRG), and spotted fever group (SRG). Nodes are colored by posterior probability. (B) Field sites relative to the American robin and dark-eyed junco distribution with the estimated breeding origins of the two PCR-positive birds. Geographic assignments were performed using feather hydrogen of previously established known-origin juncos, robins, and other passerines (Fig. S1). Paths display median migration distance, defined as kilometers between winter capture site and median coordinates in the 90% probability breeding ground.

*Rickettsia*-positive robins and juncos were sampled in winter (12/2020 and 11/2018). Most robins wintering in Indiana remain year-round or migrate elsewhere to breed, while wintering juncos in the Appalachians include migrant and resident subspecies; we earlier determined this positive bird as migratory (*J. h. hyemalis*) (7). Using feather hydrogen isotopes and geographic assignment (Figure S1) (10), we estimated the most likely breeding site of the robin to be the Great Lakes (e.g., Wisconsin; 459 km median migration distance), whereas that of the junco ranged from the western USA to Manitoba in Canada (2,377 km median migration distance; Fig. 1B). Positive birds were thus short-or long-distance migrants and could spread their novel rickettsiae during spring migration. Further study of rickettsiae in these common birds will be important to characterize their evolutionary history, whether passerines are competent hosts, and dispersal potential in relation to bird migration.

## Acknowledgements

This work was supported in part by the Prepared for Environmental Change Grand Challenge Initiative at Indiana University, the Intelligence Community Postdoctoral Research Fellowship Program, the Virginia Society of Ornithology, the National Science Foundation (DEB-0808284, IOS-0820055, DBI-0939454), and the National Institutes of Health (T32 HD49336). We thank Janice Dispoto and Jason Weckstein for assistance with DNA extraction at Drexel University.

