## Supplementary material for "Novel *Rickettsia* spp. in two common overwintering North American passerines": Figure S1

**Figure S1.** Geographic assignments of rickettsiae-positive individuals, *Turdus migratorius* 1412-68667 (A,B) and *Junco hyemalis* 1881-71194 (C,D), based on the hydrogen isotopic value of flight feathers (for details about sample preparation, hydrogen equilibration, and isotopic analysis, see Wanamaker et al., 2020). Breeding location estimates presented in Figure 1 in the main text (B,D) reflect cells within the top 10% highest posterior probability based on Bayesian assignment (A,C). Geographic assignments were performed in R using the *assignR* package (Ma et al., 2020), using growing-season-precipitation (GSP; Bowen et al., 2005) and guild-specific (short-distance migrant, non-ground foraging) and species-specific resident calibration datasets for *T. migratorius* and *J. hyemalis*, respectively.  The calibration model for *T. migratorius* produced an intercept of -39.9, approximating the diet-to-tissue discrimination factor, and explained 69% of the isotopic variation in resident individuals.  The calibration models for *J. hyemalis* produced an intercept of -21.3, approximating the diet-to-tissue discrimination factor, and explained 48% of the isotopic variation in resident individuals. Seasonal ranges from IUCN (2019) are shown in light gray underneath estimates (species’ full range maps were used to construct geographic assignment maps).

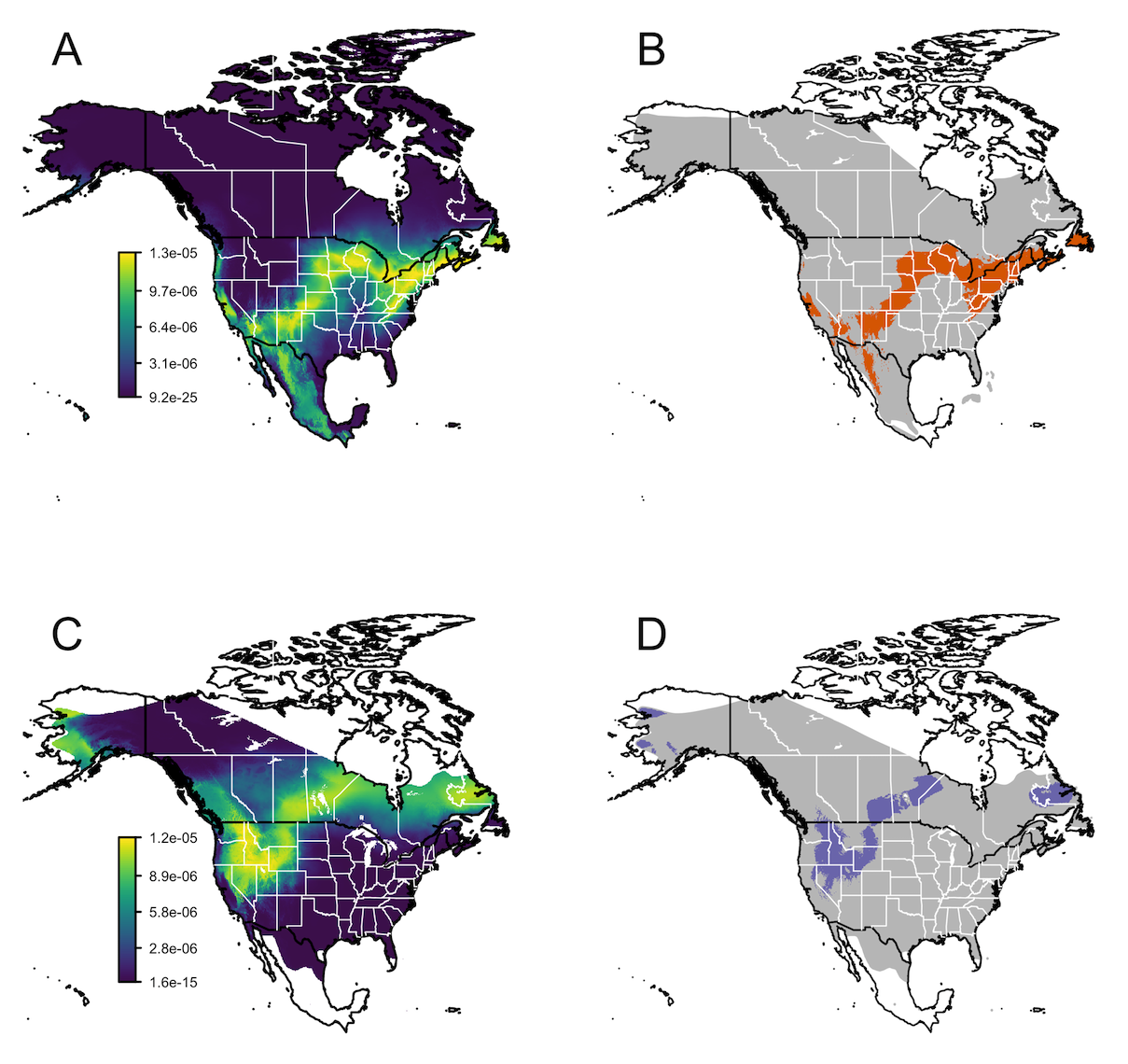
