## Supplementary material for "Novel *Rickettsia* spp. in two common overwintering North American passerines": Table S1

**Table S1.** PCR primers and amplification parameters used in this study.

| **Primers** | **Sequence** | **Organism and gene^a^** | **Product size (bp)** | **Temperature (⁰C) / time (seconds)** | | | **Cycles** | **Ref** |
| --- | --- | --- | --- | --- | --- | --- | --- | --- |
|  |  |  |  | Denaturation | Annealing | Extension |  |  |
| Bart 443f | 5’-GCTATGTCTGCATTCTATCA-3’ | *Bartonella* *gltA* | 750 | 94/30 | 48/30 | 72/30 | 30 | 1 |
| Bart 1210r | 5’-GATCYTCAATCATTTCTTTCCA-3’ |  |  |  |  |  |  | 2 |
| BhCS 781p | 5’-GGGGACCAGCTCATGGTGG-3’ |  | 370 | 94/30 | 55/30 | 72/30 | 30 | 3 |
| BhCS 1137n | 5’-AATGCAAAAAGAACAGTAAACA-3’ |  |  |  |  |  |  | 3 |
| BSP 16S1A | 5’-CTAACGCTGGCAGTGCGTCTTAAGC-3’ | *Borrelia*  16S rRNA | 724 | 94/30 | 60/30 | 72/30 | 30 | 4 |
| BSP 16S1B | 5’-AGCGTCAGTCTTGACCCAGAAGTTC-3’ |  |  |  |  |  |  | 4 |
| BSP 16S2A | 5’-AGTCAAACGGGATGTAGCAATAC-3’ |  | ~657 | 94/30 | 55/30 | 72/30 | 30 | 4 |
| BSP 16S2B | 5’-GGTATTCTTTCTGATATCAACAG-3’ |  |  |  |  |  |  | 4 |
| RCK23/5-F | 5’-GATAGGTCRGRTGTGGAAGCAC-3’ | *Rickettsia* 23S-5S rRNA ITS | ~380 | 94/30 | 60/30 | 72/30 | 30 | 5 |
| RCK23/5-R | 5’-TCGGGAYGGGATCGTGTGTTTC-3’ |  |  |  |  |  |  | 5 |
| RCK23/5-NF | 5′-TGTGGAAGCACAGTAATGTGTG-3′ |  | ~350 | 94/30 | 55/30 | 72/30 | 30 | 6 |
| RCK23/5-NR | 5′-TCGTGTGTTTCACTCATGCT-3′ |  |  |  |  |  |  | 6 |
| UNI_16S_mycF | 5′-GGCCCATATTCCTACGGGAAGCAGCAGT-3′ | Hemotropic mycoplasma  16S rRNA | ~1000 | 95/300 | 60/60 | 72/60 | 50 | 7 |
| UNI_16S_mycR | 5′-TAGTTTGACGGGCGGTGTGTACAAGACCTG-3′ |  |  |  |  |  |  |  |

^a^*Bartonella* spp. *gltA*, citrate synthase gene; *Borrelia* spp. 16S rRNA gene; *Rickettsia* spp. 23S-5S rRNA intergenic spacer; hemotropic mycoplasma spp. 16S rRNA gene
